# Overexpression of Xanthophyll Cycle Genes Leads to Faster NPQ Acclimation in the C4 Monocot *Setaria viridis*

**DOI:** 10.1101/2024.12.23.630197

**Authors:** William D. Stone, Lucia Acosta-Gamboa, Xiaojun Kang, Feng Zhang, Lauren Owens, Li Li, Peter M. Thielen, Georg Jander, Michael A. Gore, Thomas J. Lawton

## Abstract

Slow relaxation and induction of Non-Photochemical Quenching (NPQ) limits plant productivity under fluctuating light in C3 plants but it is unknown if this trait extends to C4 plants. Three genes have been implicated in determining NPQ induction and relaxation rates under intermittent leaf shading; *VDE* and *ZEP* interconvert the xanthophyll pigments violaxanthin and zeaxanthin, and PsbS regulates the overall level of NPQ. Here we report the overexpression of the *Arabidopsis thaliana VDE, ZEP*, and *PsbS* in the NADP-Malic Enzyme C4 monocot *Setaria viridis*. We demonstrate these transgenic plants have faster NPQ induction and relaxation under fluctuating light but observe no impact on Photosystem II photochemistry yield. Under illumination, faster NPQ induction resulted in higher transient NPQ levels and a lower overall chlorophyll fluorescence yield. Together this introduces the potential for lower productivity due to over-quenching of chlorophyll during light fluctuation. Although we observed faster NPQ acclimation rates in our transgenic lines, hyperspectral imaging and pigment analysis suggest that there is little photosynthetic advantage to increasing NPQ acclimation over wildtype due to naturally fast C4 acclimation rates. These results indicate that further work is necessary in evaluating the role of NPQ acclimation in C4 growth dynamics towards the goal of improving crop productivity.

## Introduction

Non-Photochemical Quenching (NPQ) is composed of a number of distinct mechanisms plants utilize to safely dissipate excess light, avoid photosystem damage, and prevent reactive oxygen species (ROS) production. NPQ is typically divided into four mechanisms with varying time responses. qE (energy dependent quenching) acts on a seconds timescale in response to an excess pH gradient across thylakoid membranes (Baker, 2008). qZ (zeaxanthin dependent) acts on a minutes timescale to remodel photosystems through the binding of zeaxanthin to light harvesting complexes and enhance energy dissipation (Nilkens et al., 2010). qT (state transitions) operates on a minutes timescale and to remodel LHC architecture via phosphorylation is typically only relevant at low light intensities (Baker, 2008). Finally, qI (photoinhibition) acts over hours to days to respond to consistently excessive light levels by modifying photosystem II as a long term stress response (Long et al., 1994). The exact regulation and implementation of these mechanisms remains actively investigated, but three genes have been implicated in qE and qZ regulation: Violaxanthin De-Epoxidase (*VDE*) and Zeaxanthin Epoxidase (*ZEP*), which interconvert violaxanthin and its quenching form zeaxanthin. Additionally, the Photosystem II subunit PsbS has been implicated as a key regulator of NPQ affecting NPQ levels through direct sensing of the thylakoid pH gradient (Kiss et al., 2008; Li et al., 2002). *PsbS* expression has been correlated with the overall amount of both transient and steady state qE component of NPQ.

Under field conditions, light levels are dynamic due to transient shading of leaves by clouds and other plants. To achieve maximum productivity, plants must dynamically modulate their levels of NPQ to match incident irradiance. In C3 plants, the slow relaxation rate of the qZ components of NPQ reduces productivity by diverting energy away from photosynthesis (Murchie and Niyogi, 2011). In a seminal field study of transgenic tobacco, overexpression of *PsbS, VDE*, and *ZEP* increased productivity and NPQ induction and relaxation rates (Kromdijk et al., 2016). This was extended to soybean, a C3 crop, and increased seed yield was realized without impacting seed quality (De Souza et al., 2022). Further work in *Arabidopsis thaliana* (Arabidopsis) and potato have replicated the increase in NPQ acclimation rates but indicated that productivity advances may be species or growth condition dependent (Garcia-Molina and Leister, 2020; Lehretz et al., 2022).The extensibility of this hypothesis to C4 plants is not well understood and merits further exploration (Sales et al., 2021). Little information is available about the differences, if any, between C4 and C3 NPQ mechanisms. However, C4 photosynthesis is distinguishable by its spatial separation of the photosynthetic light reaction and carbon fixation into different cell types, along with an increased prevalence of Photosystem I (PSI), cyclic electron transport, and a higher ATP requirement. While C4 use under fluctuating light less efficiently than C3 plants, it has recently been hypothesized that the slow speed of dark side reactions rather than NPQ limit productivity and in fact C4 plants naturally induce NPQ faster that C3 plants (Li et al., 2023; Wang et al., 2021). In our study, we investigate whether NPQ acclimation rates in C4 plants, specifically *Setaria viridis* (Setaria), can be further increased through the overexpression of Arabidopsis *PsbS, VDE*, and *ZEP*.

## Results

To test the impact of *PsbS, VDE*, and *ZEP* overexpression on NPQ in NADP-ME C4 plants, we generated transgenic Setaria lines overexpressing the Arabidopsis homologs of these genes (Brutnell et al., 2010). Homozygous, single insertion T_3_ lines were identified for three unique transformation events utilizing Oxford Nanopore sequencing. Three high expressing lines were identified by RT-qPCR on the T_1_ generation and sequencing was performed on the T_2_ generation (data not shown). Computational analysis was used to identify transgenic alleles and PCR-based genotyping of T_3_ plants identified homozygous lines (supplementary data). Overexpression of the three Arabidopsis genes was confirmed by both RT-qPCR and Western blot analysis (Figure 1). Robust expression of all three transgenes was observed and the expression levels amongst the three transgenic lines was not significantly different. No impact on the expression of endogenous Setaria NPQ genes was observed, suggesting minimal impact on transcriptional regulation of endogenous genes.

**Figure 1.**
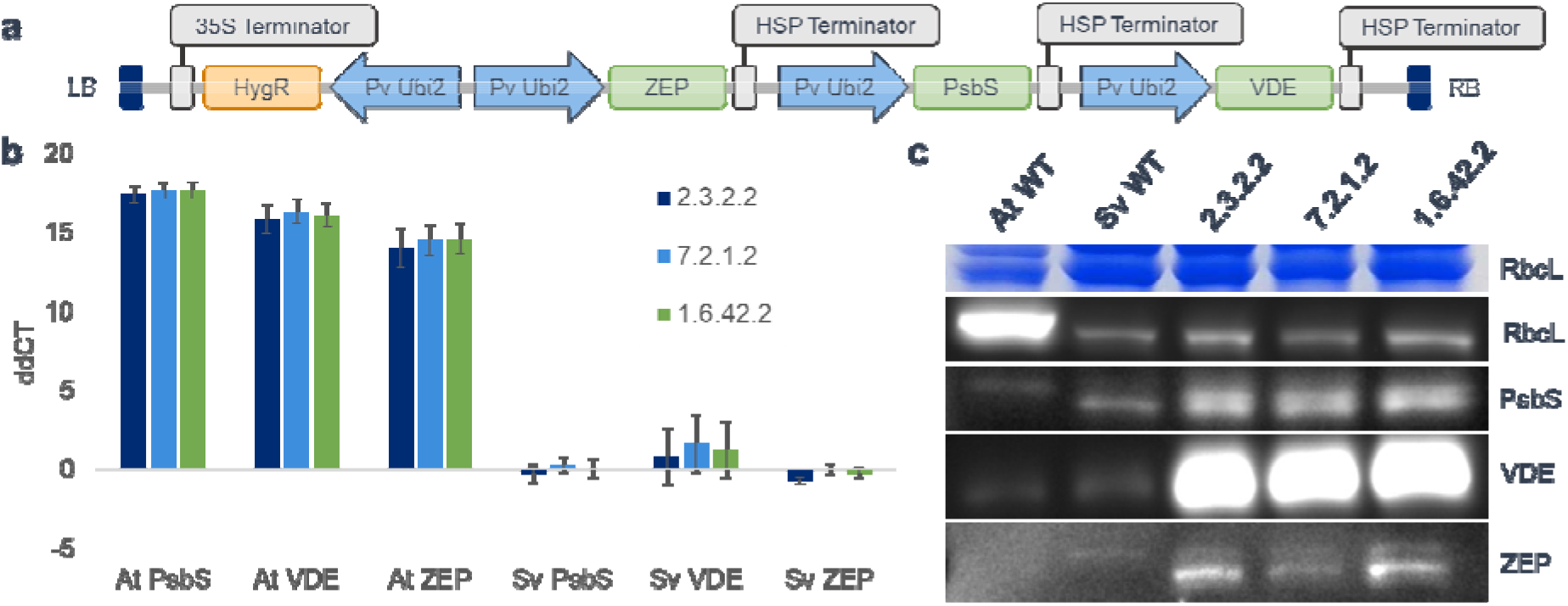
Expression of *Arabidopsis* NPQ genes in inbred T_4_ *Setaria viridis* lines. (a) diagram of overexpression construct, (b) RT-qPCR of recombinant Arabidopsis NPQ genes (At) and endogenous Setaria homologues (Sv) compared to the wildtype Setaria control. Bars are standard deviation of 3 biological replicates, (c) representative Western blot and Coomassie stain of extracted chloroplasts from each line.

Chlorophyll fluorescence was used to assess NPQ induction and relaxation dynamics in a light induction experiment (Figure 2). Dark adapted leaves were illuminated for 15 minutes with high light followed by a return to darkness for 5 minutes. No changes in maximal Photosystem II yield (Fv/FM) were observed in dark-adapted leaves indicating no long term photoinhibition (qI) in our transgenic lines. During the initial minutes of illumination, NPQ was induced more quickly and to a higher level in all three transgenic lines while no significant changes in Photosystem II yield were observed (ΦPSII). In two of the three transgenic lines, a small transient decrease in NPQ was observed one minute into the illumination regime (Figure 2f). In the third transgenic line (2.3.2.2) and the wildtype a change in the rate of NPQ induction was observed at this timepoint. Transgenic lines reached a peak NPQ value 3-4 minutes after transfer to the light, followed by a gradual decay in NPQ to a steady state value. This contrasts with the wildtype line’s monotonic increase in NPQ. During transfer from high light to darkness, NPQ relaxed faster in all 3 transgenic lines, with no impact on PSII yield observed.

**Figure 2.**
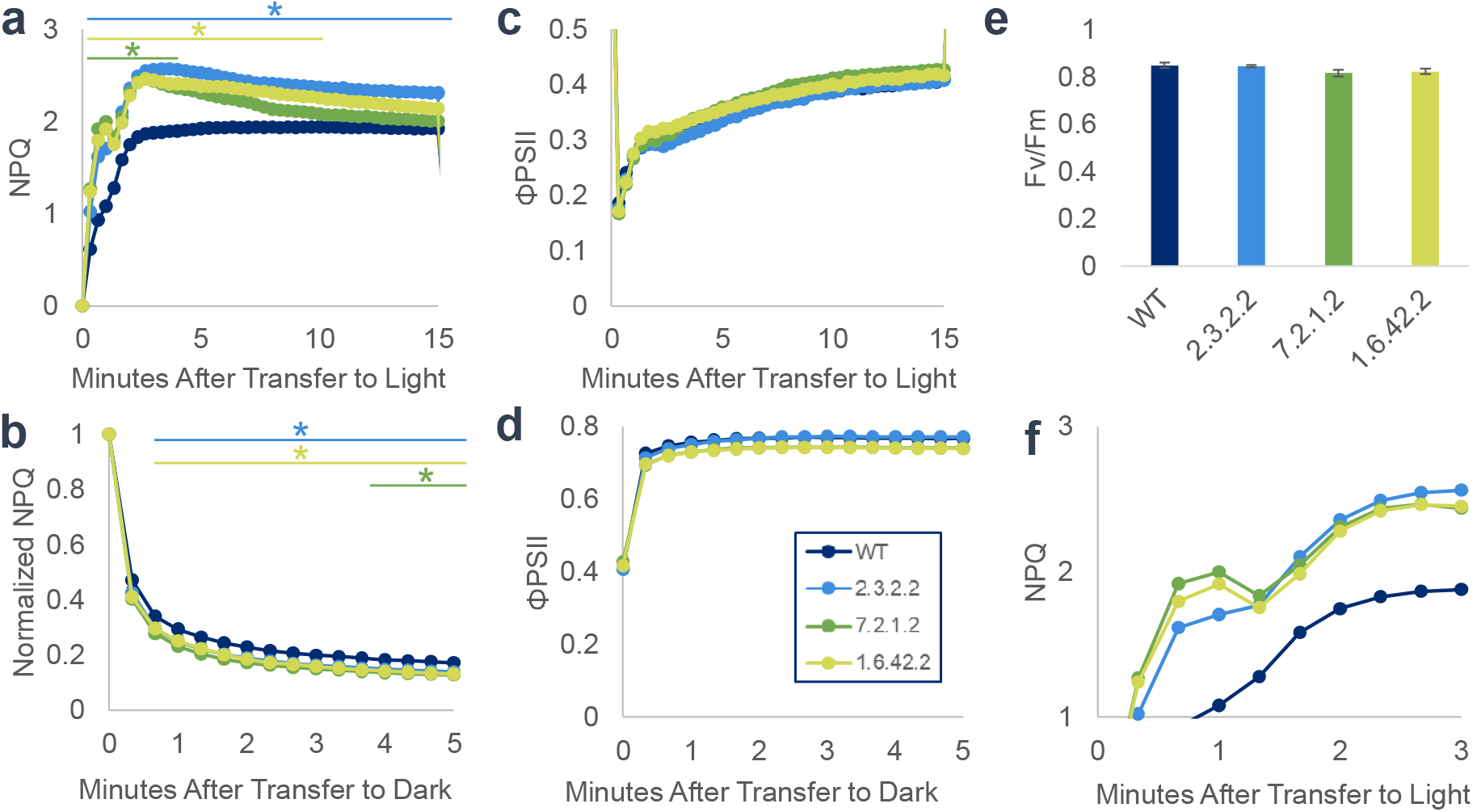
NPQ and PSII yield during photosynthesis induction and relaxation assessed by chlorophyll fluorescence. (a,b) NPQ transfer from dark to 1200 PPFD and back. (c,d) PSII yield during the same transition. (e) Dark adapted maximum PSII yield, with error bars indicating standard error. (f) Detail of a. Data represent averages of 8 biological replicates. Statistical significance between wildtype and transgenic lines is denoted by bars and asterisks (two sided t test, p<0.05).

When illuminated, NPQ was induced to a higher level in transgenic lines, without a corresponding decrease in PSII yield. This suggests a higher degree of chlorophyll quenching compared to the wildtype. To confirm this, we examined transient and steady state chlorophyll fluorescence yields (Fs) under illumination. Chlorophyll fluorescence yields were reduced compared to the wildtype under illumination in all transgenic lines, but this difference rapidly disappeared when transferred to the dark (figure 3a). To determine if this result scales across an entire plant canopy, we utilized high temporal resolution VISNIR hyperspectral imaging. We measured the Photochemical Reflectance Index (PRI), a measure positively correlated with NPQ, of individual plant canopies during a transition from low to higher light (Figure 3b) (Sukhova and Sukhov 2018). In two of three transgenic lines, PRI was substantially higher than in the wildtype under both illumination conditions. These data also confirmed the temporal dynamics of NPQ induction observed with chlorophyll fluorescence. Under moderate light intensity change, all plants reached a maximal value of the PRI within one minute of transfer.

**Figure 3.**
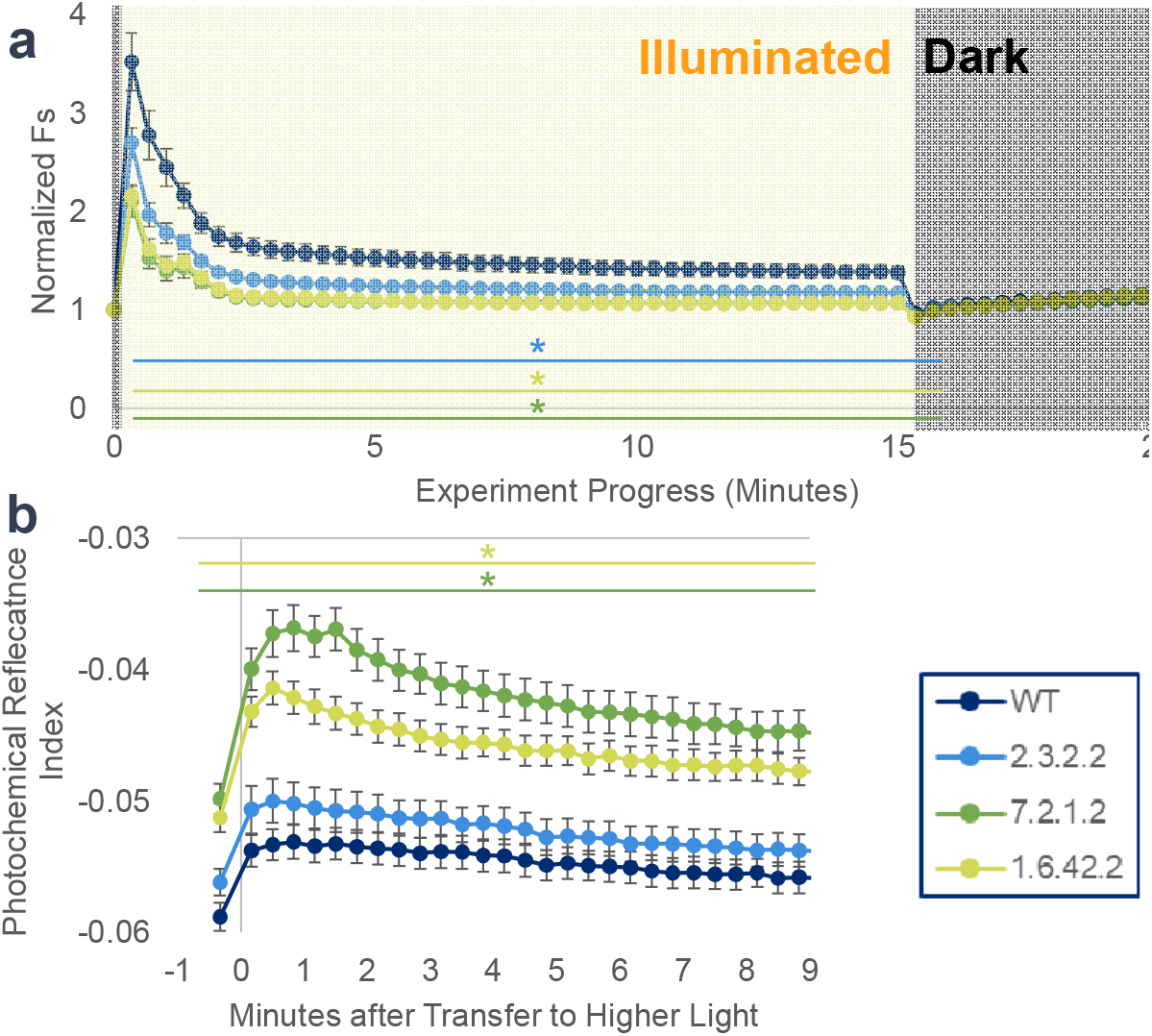
Timeseries Fluorescence and HSI dynamics (a) Dark normalized chlorophyll fluorescence yield during photosynthesis induction and relaxation. Leaves were dark adapted at start, then, illuminated for 15 minutes at 1200 PPFD, and finally returned to darkness for 5 minutes. (b) Photochemical Reflectance Index of canopies during transfer from 150 to 500 PPFD, as assessed by hyperspectral imaging. Data represent averages of 8 biological replicates, with error bars indicating standard errors. Statistical significance between wildtype and transgenic lines is denoted by bars and asterisks (two sided t test, p<.05).

Pigment analysis was performed using HPLC of lyophilized leaf tissues to further confirm NPQ dynamics. In mature leaves, a decrease in total chlorophyll content was observed in the transgenic lines (Figure 4a). Violaxanthin content was measured as a proxy for xanthophyll de-epoxidation under constant and variable light regimes (Figure 4b). Violaxanthin content was lower than of the wildtype under all illuminated conditions and in 2 out of 3 dark-adapted lines. When subjected to four cycles of alternating 200 and 1200 PPFD light, there was no difference in violaxanthin content compared to plants adapted to the final step (1200 PPFD), regardless of genotype.

**Figure 4.**
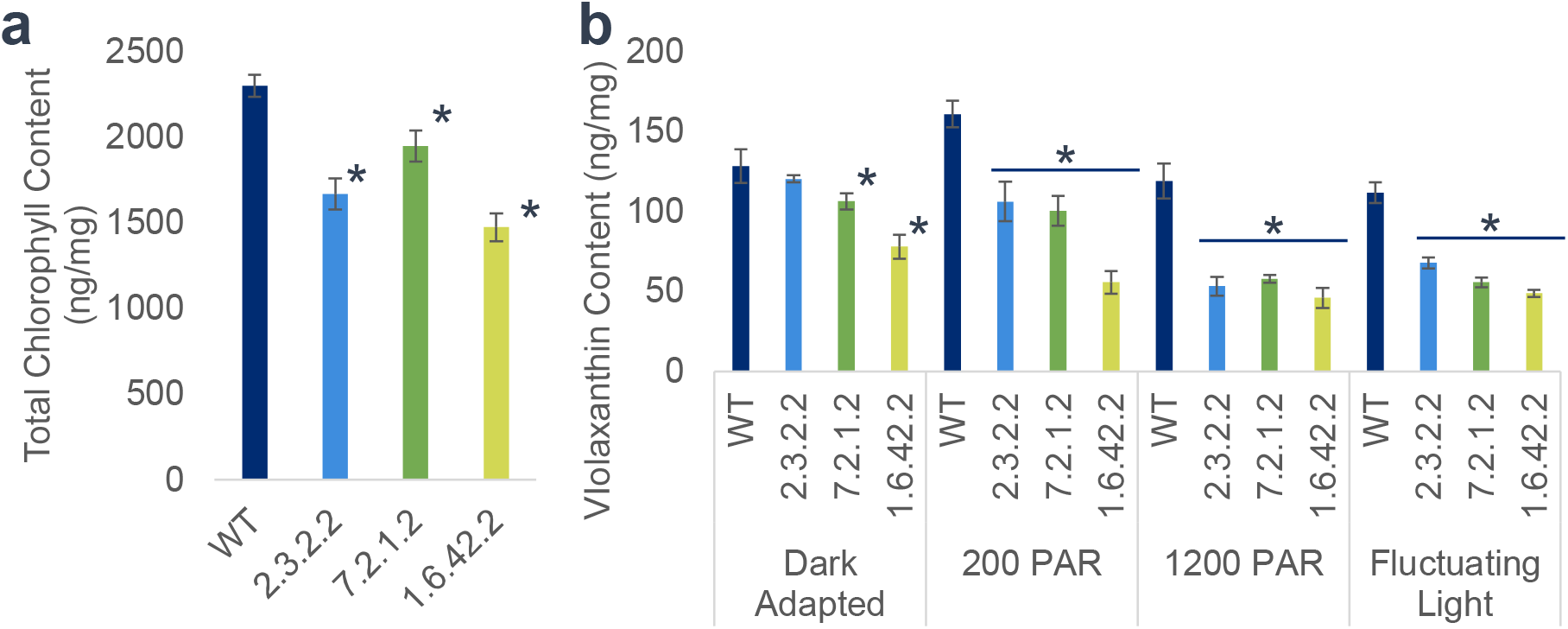
Pigment Dynamics. (a) Total chlorophyll content of mature leaves assessed by HPLC. Data represent the average of 16 biological replicates, and error bars indicate standard errors. (b) Violaxanthin content of dark-adapted, 200 PPFD adapted, 1200 PPFD adapted, and fluctuating light adapted leaves assessed by HPLC. Data represent the average of 4 biological replicates and error bars indicate standard errors. The content of each pigment is normalized to lyophilized leaf mass. Statistical significance is denoted by bars and asterisks (two sided t test, p<0.05).

## Discussion

The overexpression of *PsbS, VDE*, and *ZEP* resulted in the desired phenotype of increased NPQ acclimation and relaxation rates. However, several unexpected phenotypes were also observed. During induction, engineered lines achieved a NPQ induction rate approaching the steady state rate approximately one minute after transfer to the light rather than over two minutes in the wildtype Setaria. However, in engineered lines NPQ unexpectedly continued to climb overshooting the steady state rate before decaying to steady state, a feature not seen in wildtype. During relaxation from steady state illumination, all lines demonstrated a monotonic decrease in NPQ with engineered lines having a faster rate of decay.

In our transgenic lines, we observed two NPQ induction and relaxation events, distinguishable by their temporal response during light induction. First, there was a rapid increase in NPQ followed by a transient decrease around one minute after transfer to light (Figure 2f). Given the timing of this feature, it can be attributed to the qE component of NPQ. In our transgenic lines, qE acclimation was both accelerated and saturates at a higher level. After induction to approximately the steady state NPQ value, qE appears to decay rapidly resulting in a transient decrease in NPQ in two out of three lines. This feature was not observed in similar studies in C3 tobacco and soybean, suggesting that qE may play an altered role in C4 NPQ (De Souza et al., 2022; Kromdijk et al., 2016). Whether this feature is due to an increased qE capacity or simply enhanced qE activity in our transgenic lines is unknown. However, studies in C3 plants have correlated *PsbS* expression levels with the qE capacity, suggesting that this elevated capacity is due to overexpression of this gene (Kiss et al., 2008). Mechanistically, the role of zeaxanthin in *PsbS* mediated qE is debated with some studies implicating it and others concluding *PsbS* induces NPQ solely in response to thylakoid pH gradient (Nicol and Croce, 2021; Lee et al 2024).

A second distinct NPQ induction event in our transgenic lines was observed in the subsequent increase in NPQ to a peak value 3-5 minutes after illumination. Total NPQ overshoots and subsequently declines to a steady-state value on a minutes timescale. Given the timing, this feature can be attributed to the qZ component of NPQ. This feature was observed in C3 plants and has been observed in C3 lines engineered to overexpress these three genes (Kromdijk et al., 2016). Interestingly, overshoot of qZ requires greater than 10 minutes to relax to a steady state value. The rate of this acclimation has been speculated to be determined to the slow association/dissociation kinetics of xanthophyll pigment binding to specific sites on LHCII and qZ is determined by the xanthophyll de-epoxidation state with a lag (Nilkens et al., 2010). We hypothesize that the qZ overshoot in our engineered lines is a reflection of increased de-epoxidation during early light induction driven by VDE overexpression.

The decay of NPQ from its peak ∼3 minutes after transfer to light in our transgenic lines complements the slow induction of photochemistry (as measured as ΦPSII) and results in a consistently reduced fluorescence yield (Fs) under illumination (Figure 3). This suggests that chlorophyll fluorescence is more effectively quenched during this transient increase in light, thereby avoiding ROS production and damage. A potential benefit of this increased quenching could be increased stress tolerance. A *PsbS* overexpressing tobacco line with similar increases in transient NPQ demonstrated increased water-use efficiency and drought tolerance (Głowacka et al., 2018). Further studies may elucidate the impact of each of these genes on stress incidence (eg. drought, light, and temperature) in our generated Setaria transgenic lines, as C4 acclimation to stress differs from C3 plants (Anderson et al., 2021).

One limitation of this study is the sole use of chlorophyll fluorescence to measure photosynthesis. No changes in Photosystem II yield were observed in this study, suggesting but not proving there was no impact on photosynthetic productivity. CO_2_ gas exchange measurements could potentially show variations in CO_2_ fixation rates due to the spatial separation of C4 metabolism, especially under fluctuating light conditions. One possible point of comparison between C3 and C4 plants is the degree of violaxanthin epoxidation under fluctuating light vs steady acclimation. Kromdijk et al. found that three minutes of induction was insufficient for wildtype tobacco to achieve a xanthophyll epoxidation state consistent with the higher light level, but was sufficient for engineered lines expressing *PsbS, VDE*, and *ZEP* (Kromdijk et al., 2016). Although we observe marginally faster NPQ relaxation rates in our Setaria transgenic lines, we find no lack of agreement in vioxanthin content between wildtype plants treated with fluctuating light for 3 minutes and plants adapted to the final light step (1200 PPFD) (Figure 4b). Additionally, our timeseries hyperspectral imaging (Figure 3b) revealed that the wildtype line increased NPQ in response to a moderate shift in illumination on a timescale of around one minute. This observation was consistent in our engineered lines and aligns with observations from other C4 species where faster relaxation was observed(Wang et al., 2021). Taken together, this suggests the potential gains in photosynthetic productivity through faster NPQ acclimation in C4 plants are less significant than C3 plants because C4 are already well adapted to fluctuating light.

Future work with our engineered Setaria lines should include growth studies to determine their productivity and stress tolerance lines under dynamic conditions. Further tuning of NPQ gene expression through the use of alternative promoters or terminators, or single gene expression constructs may also prove fruitful understanding the mechanisms of C4 NPQ and their regulation as well as achieving elevated productivity.

## Materials and Methods

### Plant Culture

Unless otherwise noted, all plants were grown in controlled environment chambers (Conviron) with 250 μmol m^-2^ s^-1 PPFD^ and 25° C days (16 hours)/23° C nights (8 hours) in a mixture of All-Purpose Potting Mix (Promix) and Carolina Potting Soil (Carolina Biological Supply). To achieve high germination rates, seeds were soaked in a 5% Liquid Smoke solution (Wright) at 30° C for 48 hours or the seed coat was removed by agitation.

### Generation of Inbred transgenic plants

The plasmid contained coding sequences of three genes from *A. thaliana*: violaxanthin de-epoxidase (*AtVDE*, At1G08550.1), Photosystem II subunit S (*AtPsbS*, AT1G44575.1) and zeaxanthin epoxidase (*AtZEP*, AT5G67030.1), previously confirmed to exhibit activity in *N. benthamiana* (Kromdijk et al., 2016). The coding sequence of each gene was synthesized by Integrated DNA Technologies, Inc, optimized with *Setaria viridis* codons. The individual coding sequences were cloned into three constructs (pMOD_A3013, Addgene #91046; pMOD_B2112, Addgene #91064; pMOD_C3013; Addgene #91100) between the *PvUbi2* promoter and terminator using the restriction enzymes, *SbfI* and *XhoI*, respectively. The resulting constructs were then assembled into a destination T-DNA vector (pTRANS_253, Addgene #91125) using the Golden Gate cloning method (Čermák et al., 2017). The final T-DNA construct, pMG121, was used for Setaria transformation.

Transgenic *Setaria viridis MEO34v* were generated by the Boyce Thompson Institute Center for Plant Biotechnology Research using the Agrobacterium mediated callus transformation method (Van Eck and Swartwood, 2015). At the T_1_ generation, lines were screened for transgene expression by RT-qPCR (data not shown). The three highest expressing lines were advanced to the T_2_ generation and multiple plants were PCR screened for the presence of hygromycin. For each transgenic line, leaf tissue was pooled from more than 5 PCR positive plants for genomic DNA extraction. Briefly, highly intact genomic DNA was extracted using CTAB buffer followed with phenol-chloroform separation following a 48 hour dark treatment to deplete leaf carbohydrates (Thielen et al., 2020). DNA was sequenced using Oxford Nanopore and transgene insertions were identified utilizing Transgene Insertion Finder (TgIF; https://github.com/jhuapl-bio/TgIF).

TgIF identified one transgenic insertion for the 2.3.2 and 7.2.1 lines and two insertions for the 1.6.42. The 1.6.42 line was bred to achieve a homozygous single insertion line in the in the T3 generation. Each transgenic allele was confirmed by PCR-based genotyping with primers sets designed approximately 100 bp upstream and downstream of each of the insertion loci. The number of insertions present in each line was confirmed using Mendelian inheritance ratios of the hygromycin cassette in the T_3_ generation, as assessed by PCR-based genotyping (data not shown).

### PCR-Based Genotyping

Genotyping was conducted utilizing the Plant Phire Direct Polymerase (Thermo Fisher Scientific). Briefly, a 2mm biopsy punch was taken for each plant and placed in 20 µl of Phire DNA dilution buffer. The leaf sample was crushed with a sterile pipette tip and 1 µl of the resulting supernatant was used as template for the PCR assay. To identify homozygous single transgene insertion plants, T_3_ plants for each line were first screened first for the presence of the hygromycin cassette and secondly for the presence of transgenic alleles utilizing a three primer allele discrimination PCR assay. The three primers utilized were: the hygromycin reverse primer, and the two genomic locus specific primers. Transgenic alleles yielded a ∼900bp PCR product, while wildtype alleles yield a 250bp PCR product. Heterozygous plants yielded both products. Primer sequences and genotyping gel electropherograms maybe found in the supplementary data.

### RTqPCR

Total RNA was extracted from a single leaf of 3-week-old T_4_ Setaria lines using the Directzol RNA kit (Zymo) including on-column DNAse treatment. cDNA was reverse transcribed with the Maxima H Minus First Strand cDNA Synthesis Kit (Thermo Scientific). RTqPCR was performed using the PowerUp SYBR Green Master Mix (Applied Biosystems) using a two-step protocol: 2 minutes at 50C, then 2 minutes at 95°C, then 40 cycles of 15 seconds at 95°C and 1 minute at 61°C. Transcripts for both the host and transgenic homologs of *PsbS, VDE*, and *ZEP* were quantified and normalized to an internal reference gene (Cullin, Si021373) (Lambret-Frotté et al., 2015).

### Western Blot

Total Leaf chloroplast protein was extracted from the most recent fully developed leaf of 3-week-old Setaria plants following 2-day dark treatment to deplete chloroplast carbohydrate content. Intact chloroplasts were extracted following published methods (Chen et al., 2016). Briefly approximately 3 grams of leaves were pooled from 3 sibling plants and ground in a mortar and pestle with ice-cold extraction buffer. Chloroplasts were isolated by filtering the lysate twice with Miracloth (Millipore Sigma), followed by pelleting twice by spinning at 2,000g for 5 minutes. The resulting chloroplast pellet was resuspended and the chlorophyll content quantified spectrophotometrically (Porra et al., 1989).

10 µg of total chlorophyll was diluted into Laemmli buffer and fractionated on 4-20% Tris-glycine SDS-PAGE gels. Gels were semi-dry transferred to PVDF membranes and then blocked with 5% non-fat milk in Tris-buffered saline with Tween (TBST) (iblot 2, Thermo Fisher Scientific). All antibodies were diluted in 1:2,000 in TBST with 5% milk. The primary antibodies used were: *RbcL* (PhytoAB, PHY0346), PsbS (Agrisera, AS09533), *VDE* (Agrisera, AS153091), and ZEP (AS153092). The secondary antibody used was conjugated to Alexa Fluor 488 (Abcam, ab150077).

### Chlorophyll Fluorescence

Chlorophyll fluorescence measurements were conducted using a MultispeQ fluorometer (PhotosynQ). Approximately 3-week-old T_4_ plants were dark adapted for 20 minutes before every measurement. Fluorescence experiments consisted of a single dark-adapted measurement, then 15 minutes 1200 PPFD illumination and followed by 5 minutes of darkness. Measurements were taken every 20 seconds over the course of the experiment and consisted of a variable fluorescence yield measurement (Fo or Fs), followed by a maximum fluorescence yield measurement (Fm or Fm’) following a 400ms pulse of saturating light (15000 PPFD). Fluorescence yield was induced by a 100ms 400 PPFD measurement pulse.

Maximal Photosystem II efficiency was calculated as: Fv/Fm = (Fm-Fo)/Fm

Photosystem II yield was calculated as (Fm’-Fs)/Fm’

NPQ was calculated according to the Stern-Volmer model(Bilger and Björkman, 1994): Fm/FM’-1

### Hyperspectral Imaging

Timeseries HSI was conducted with a pushbroom VISNIR (400-1000nm) sensor (SOC710, Surface optics) on 3-week-old T_4_ plants adapted to 150 PPFD for 15 minutes. After the initial HSI collection illumination was increased to approximately 500 PPFD, and images were collected approximately every 20 seconds for 9.5 minutes. Reflectance was normalized to a Spectralon PTFE reflectance standard (Ocean Optics) included in each image. Images were analyzed in MATLAB (R2020b, Mathworks) using a custom GUI. Briefly, leaf tissue was segmented by filtering pixels with a Modified Chlorophyll Absorption Reflectance Index 2 value greater than 0.6. A region of interest was manually specified to include the entire plant canopy. Hyperspectral indices were calculated using the following formula:

MCARI2 (Haboudane et al., 2004): (1.5(2.5(ρ800-ρ670)-1.3(ρ800-ρ550)))/(√((2_*_ρ800+1)^2^-(6_*_ρ800-5_*_√ρ670)-.5))

PRI (PEÑUELAS et al., 1995): (ρ570-ρ531)/(ρ531+ρ570)

### HPLC

Three-week-old T_4_ plants were adapted to dark, 200 PPFD, or 1500 PPFD for at least 15 minutes before harvesting. Fluctuating light samples were subjected to 4 cycles of 200->1500 PPFD transitions with 3 minutes at each level. At harvest, the most recent fully developed leaf was flash frozen in liquid nitrogen and homogenized with a mortar and pestle.

Carotenoid and chlorophyll pigments from the ground leaf tissue were extracted and analyzed according to previously described methods (Yazdani et al., 2019). Briefly, leaf samples (350 μg) were ground with 0.8 mL of 80% acetone and mixed with 0.4 mL of ethyl acetate and 0.4 mL of water. Following centrifugation, the upper pigment-containing phase was dried and resuspended. The extracted pigments were analyzed using an Acquity UPC2 system (Waters). Quantification of chlorophyll and individual carotenoids was achieved by comparing peak areas to the calibration curves constructed with commercial standards.

## Acknowledgements

We would like to thank Dr. Joyce Van Eck and the Boyce Thompson Institute Center for Plant Biotechnology Research for generating the transgenic plants and their assistance with growth protocols. We also extend our gratitude to Dr. Ryokei Tanaka for his support in conducting statistical analysis for T_2_ plant screening, and Dr. Luke Gregory for providing comments on an earlier version of the manuscript.

## Author Contributions

T.J.L and M.A.G. Conceived the research plan and obtained funding. W.D.S performed expression, fluorescence, and HSI experiments. X.K. performed cloning and design under supervision of F.Z, L.A.G performed preliminary expression analysis and generation advancement, and L.A.G., G.J., and M.A.G. interpreted early generation expression and genotyping results. L.L and L.O performed HPLC of carotenoid pigments. W.D.S., T.J.L and M.A.G prepared the manuscript.

## Data Availability

Data presented in this study are available in the supplemental section and from the corresponding author upon request.

