## Supplementary Materials for "Overexpression of Xanthophyll Cycle Genes Leads to Faster NPQ Acclimation in the C4 Monocot *Setaria viridis*"

**Supplementary Data – Table of Contents**

Supplementary Table 1. DNA Primer sequences

Supplementary Table 2. TGIF identified transgenic insertion sites by line

Supplementary Figure 1. Genotyping PCR of the 2.3.2 line

Supplementary Figure 2. Genotyping PCR of the 1.6.42 line

Supplementary Figure 3. Genotyping PCR of the 7.2.1 line

**Supplementary Data**

**Supplementary Table 1**. DNA Primer sequences

| Purpose | Primer Name | Significance | Sequence |
| --- | --- | --- | --- |
| RT-qPCR | WDS-159 | At_PsbS_F | GGTATTGCGTTCTCGCTTAT |
|  | WDS-160 | At_PsbS-R | GGTAATGAACTTCCCGTTCC |
|  | WDS-161 | At_VDE_F | AGAAGGCCGCTAAGTCTAT |
|  | WDS-162 | At_VDE_R | AGAGTCATCTCAGTCCTACC |
|  | WDS-163 | At_ZEP_F | CGGGTGATCTACAAAGATGG |
|  | WDS-164 | At_ZEP_R | TCGGAGTCTTGCGGATAA |
|  | WDS-242 | Sv_psbs_F | TGAACAGGCACTCCAAGTC |
|  | WDS-243 | Sv_psbs_R | GAAGATACCGTCCTCAACCTTC |
|  | WDS-246 | Sv_vde_F | GGCATGTGTAGTTCTGGTAGTC |
|  | WDS-247 | Sv_vde_R | TGCATTTGGCCAGCTCTAT |
|  | WDS-250 | Sv_zep_F | CCCAGTCACAAGGGTCATTAG |
|  | WDS-251 | Sv_zep_R | GACTACGTGGCTACCATTCAATA |
|  | Sv_Cullin_F | Cullin_F (Housekeeping) | TATGGGTCATCAACAGCTTGTC |
|  | Sv_Cullin_R | Cullin_R (Housekeeping) | GTAGTCCCTCGTGATGAGATCC |
| Genotyping PCR | WDS-139 | Hygromycin_R | CTCTCGATGAGCTGATGCTTTG |
|  | WDS-140 | Hygromycin_F | GCTGTTATGCGGCCATTGT |
|  | WDS-167 | 2.3.2_Chr2_F | GTATATAATGGAGTACGTGCATTGG |
|  | WDS-168 | 2.3.2_Chr2_R | AAAGCAAGATTCGACTACGTAC |
|  | WDS-169 | 1.5.16_Chr1_F | CGTGTTCTCCAGCTTCAAG |
|  | WDS-170 | 1.5.16_Chr1_R | CACTCGTTTCCGCGAATTG |
|  | WDS-173 | 1.5.16_Chr4_F | GCACCTTCTTCGTCTATCAC |
|  | WDS-174 | 1.5.16_Chr4_R | CTTGGTCTTGCTGATAATCTGC |
|  | WDS-238 | 7.2.1_Chr5_F | GCAGAAACTACAACTCTGGAC |
|  | WDS-239 | 7.2.1_Chr5_R | CAAATTGGCAAAATTGGCAAC |

**Supplementary Table 2.** TGIF identified transgenic insertion sites by line

| Transgenic Line | Chromosome | Insertion Site |
| --- | --- | --- |
| 2-3-2 | Chr 2 | 2692491 -2692532 |
| 1-6-42 | Chr 1 | 27376900 -27376955 |
|  | Chr 4 | 13709748 -13709773 |
| 7-2-1 | Chr 5 | 9957322 - 9957344 |

**Supplementary Figure 1.** Genotyping PCR of the 2.3.2 line

Agarose gels of representative genotyping PCR of the T3 2.3.2 line. (A) hygromycin primer set, (b) Chromosome 2 T-DNA insertion primer set. From left: lane 1 ladder, lane 2 wildtype, lane 3 heterozygous T3 sibling, lane 4 homozygous 2.3.2 line.

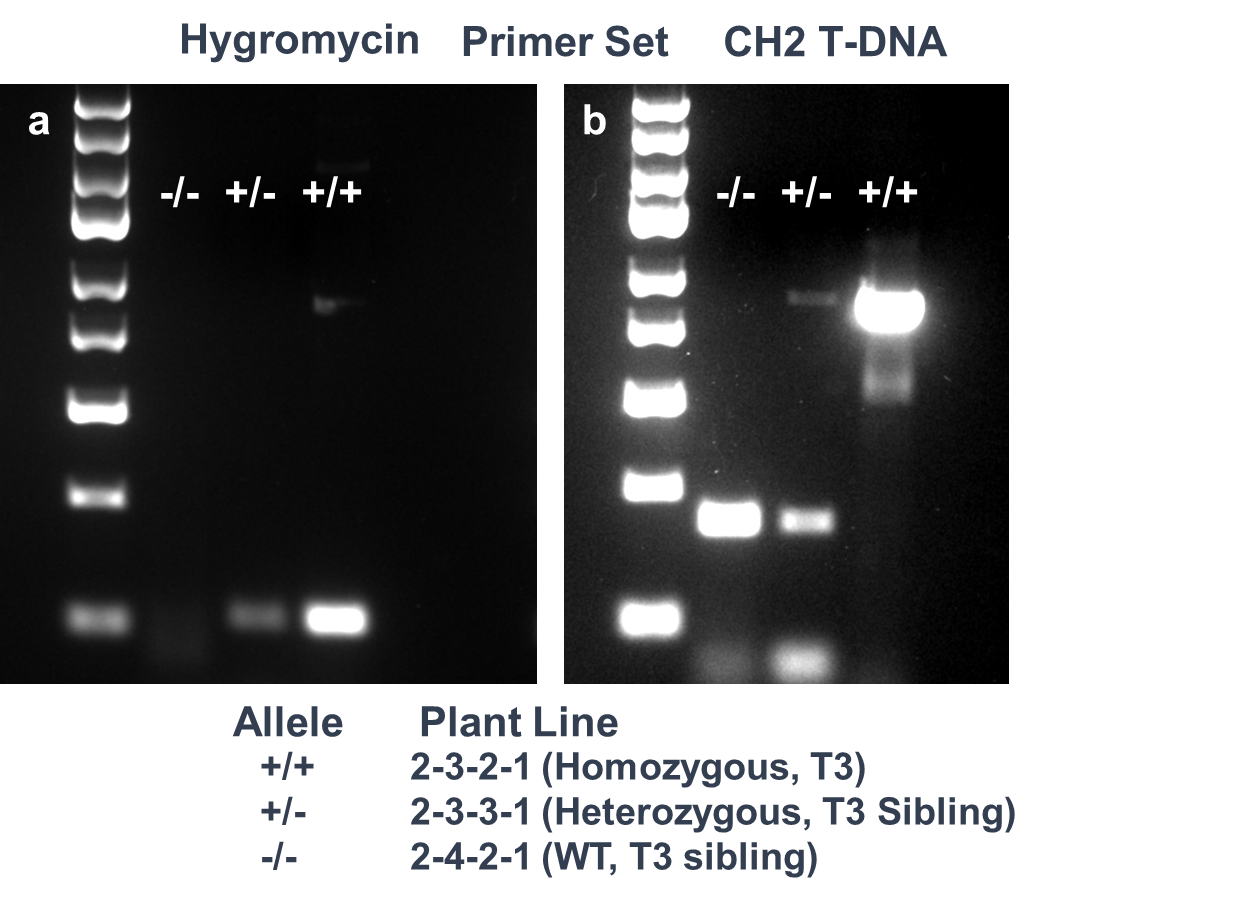

**Supplementary Figure 2.** Genotyping PCR of the 1.6.42 line

Agarose gels of representative genotyping PCR of the T3 1.6.42 line. (A) hygromycin primer set, (b) Chromosome 1 T-DNA insertion primer set, (c) Chromosome 4 T-DNA insertion primer set. From left: lane 1 ladder, lane 2 wildtype, lane 3 heterozygous T3 sibling, lane 4 homozygous 1.6.42 line.

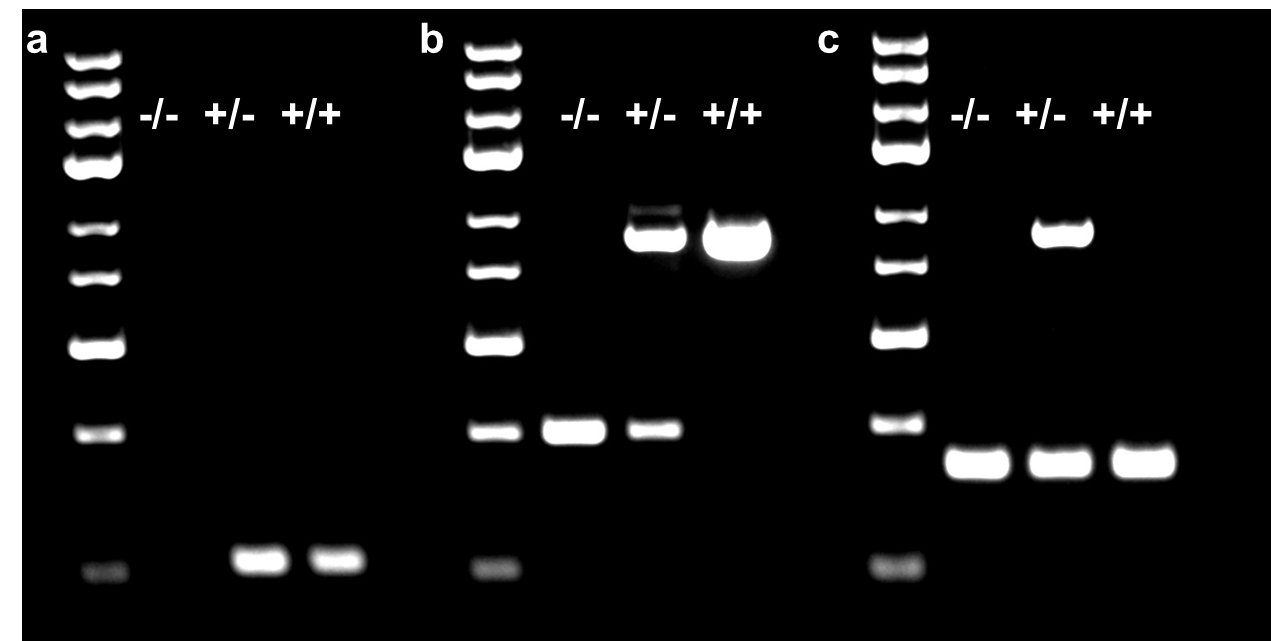

**Supplementary Figure 3.** Genotyping PCR of the 7.2.1 line

Agarose gels of representative genotyping PCR of the T3 7.2.1 line. (A) hygromycin primer set, (b) Chromosome 5 T-DNA insertion primer set. From left: lane 1 ladder, lane 2 wildtype, lane 3 heterozygous T3 sibling, lane 4 homozygous 7.2.1 line.

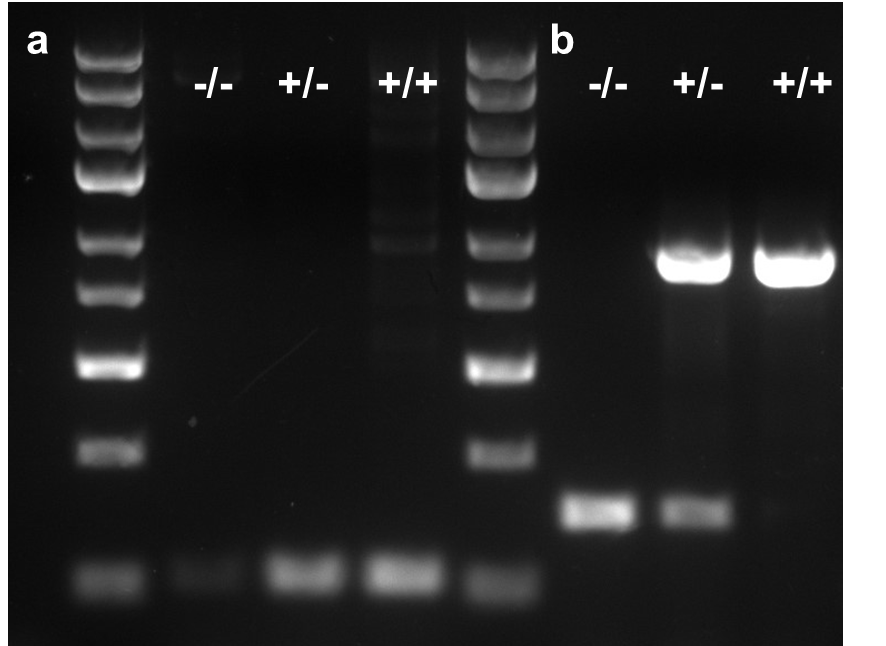
